# Isolating and Cryo-Preserving Pig Skin Cells for Single Cell RNA Sequencing Study

**DOI:** 10.1101/2021.01.31.429035

**Authors:** Li Han, Carlos P Jara, Ou Wang, Sandra Thibivilliers, Rafał K. Wóycicki, Mark A. Carlson, William H. Velander, Eliana P. Araújo, Marc Libault, Chi Zhang, Yuguo Lei

## Abstract

The Pigskin architecture and physiology are similar to these of humans. Thus, the pig model is valuable for studying skin biology and testing therapeutics for skin diseases. The single-cell RNA sequencing technology allows quantitatively analyzing cell types, cell states, signaling, and receptor-ligand interactome at single-cell resolution and at high throughput. scRNA-Seq has been used to study mouse and human skins. However, studying pigskin with scRNA-Seq is still rare. Here we described a robust method for isolating and cryo-preserving pig single cells for scRNA-Seq. We showed that pigskin could be efficiently dissociated into single cells with high cell viability using the Miltenyi Human Whole Skin Dissociation kit and the Miltenyi gentleMACS Dissociator. Also, we showed that the subsequent single cells could be cryopreserved using DMSO without causing additional cell death, cell aggregation, or changes in gene expression profiles. Using the developed protocol, we were able to identify all the major skin cell types. The protocol and results from this study will be very valuable for the skin research scientific community.

## Introduction

Annually, millions of people are affected by skin injuries and diseases^1–3^. Animal models are widely used to understand the skin physiopathology, as well as to test potential therapeutics^4,5^. Among various experimental animals, the skin architecture and physiology of pigs is one of the closest to humans^6–8^. Thus, FDA highly recommends including pigs for pre-clinical biology study and therapeutic testing^9–11^. Conventionally, the skin is investigated using low-content technologies such as the qPCR, flow cytometry, and histology. The most recently developed single-cell RNA sequencing (i.e., scRNA-Seq) technology allows simultaneously and quantitively analyze the transcriptome of thousands of individual cells. This allows new insights into the cell types, their states, signaling, receptor-ligand interactome, and their dynamics during development, disease and treatment^12–14^. Additionally, when combined with high-content immunostaining and fluorescent imaging^15^, or the cutting-edge Spatial Transcriptomics technology^12–14^, the spatial and temporal organization of these cells and their interactions can also be obtained. The scRNA-Seq has been used to study rodent^16–23^ and human^14,22,24–31^ skins. However, the study of pigskin with scRNA-Seq is still rare. and there is a critical need to develop methods for isolating high quality single cells from pigskin for scRNA-Seq.

In a typical scRNA-Seq workflow, tissues are first dissociated into single cells (i.e., the upstream single cell preparation). These freshly isolated cells are then used to prepare single cell RNA libraries mostly using the droplet-based technology (i.e., the downstream library preparation)^32^. The libraries are then sequenced using the deep sequencing technology^33^. Preparing high quality single cells is critical for the success of downstream process. Partial or incomplete tissue dissociation, cell aggregation and cell death should be avoided during the preparation. Recently studies showed that the addition of a cell preservation step between the upstream and downstream makes the scRNA-Seq workflow much more flexible, manageable, and accessible to researchers^25,34–37^. First, many animal research facilities do not have the infrastructure and expertise for preparing scRNA-Seq libraries for the fresh isolated cells. Second, most studies have multiple samples. A scRNA-Seq library preparation device allows preparing an RNA library for thousands cells per sample. However, preparing a library from multiple samples is time consuming activity, and requires multiple equipment, which is financially limiting. Third, many studies collect samples at multiple time points, and at different locations. Preparing RNA libraries at the same facilities by the same people may increase the consistency and reduce experimental variations. All these challenges can be addressed if freshly isolated single cells can be preserved. However, the preservation should not significantly change the cell viability, composition and gene expression^25,34–37^.

we report our success of isolating, cryo-preserving and scRNA-Seq of pig skin cells. We developed a robust protocol for single cell preparation and preservation. Briefly, fresh skin tissues were dissociated into single cells using the Miltenyi Biotec Whole Skin Dissociation kit and Gentle MACS Dissociator^28^. Single cells were frozen in 90% FBS+10% DMSO, stored in liquid nitrogen^34,38^. Cells were thawed, and dead cells were removed via FACS right before preparing libraries using the 10x Genomics Chromium technology. Alternatively, dead cells could be removed via MACS using the Miltenyi Dead Cell Remove Kit. Our results showed high quality single cells could be isolated and cryo-preserved with this protocol. The DMSO-based cryopreservation did not alter cell viability, composition and gene expression. We were able to identify all the major skin cells using both the fresh and frozen samples. Methods and findings from this study will be very valuable for the scientific community.

## Methods

### Harvest pig skin

Freshly euthanized healthy farm male pigs with about 30 kg of body weight were provided by the University of Nebraska-Lincoln swine facility. The dorsal skin was washed by PBS and the fur was removed entirely with a disposable scalpel and further disinfected with 70% ethanol. The full skin was harvested using sterile scissors and immediately stored in MACS Tissue Storage Solution (Miltenyi) with 1% Antibiotic-Antimycotic (Gibco). The skin was kept on ice and transported to the lab for the next steps.

### Isolate single cells from pig skin

Skin was washed with PBS, and the subcutaneous fat was scraped off using a scalpel. A full skin including the epidermis and dermis was taken with a 4 mm diameter punch (Robbins). The sample was treated with the Human Whole Skin Dissociation Kit (Miltenyi) and mechanically dissociated with the gentleMACS Dissociator (Miltenyi) following the product instructions and using the Program h_skin_01. The cell suspension was passed through a 70 μm strainer (Corning). The viability of cells was estimated by LIVE/DEAD™ Viability/Cytotoxicity Kit for mammalian cells (Invitrogen).

### Cryopreservation

The fresh isolated cell suspension was centrifuged at 300 x g for 5 minutes, 4°C. The cell pellet was resuspended at 1×10^6^ cells per ml in freeze medium containing 10% DMSO (Sigma) in 90% FBS (Gibco). 1 ml cells was put in a cryoprotective vial and placed in a pre-cold Mr. Frosty Freezing Container (Thermo Fisher) filled with isopropyl alcohol and cooled at −80°C freezer overnight before being stored in liquid N2 for long-term.

### Thawing of cells

The frozen vial was removed from the liquid N2 storage, and placed in a 37°C water bath. After thawing, the cell suspension was centrifuged at 300g for 5 min, and resuspended in DMEM + 10% FBS. The cell viability was assessed with LIVE/DEAD™ Viability/Cytotoxicity Kit and flow cytometer.

### Removing dead cells and cell aggregates using FACS

4 μL of 2 mM ethidium homodimer-1 was added to each milliliter cell suspension. cells were incubated for 20 minutes at room temperature to stain the dead cells before FACS (FACSAriaII, USA). Side-scatter and forward-scatter profiles were used to eliminate cell doublets. Living cells were gated by ethidium homodimer-1 negative. The cells were reanalyzed for purity, which typically was > 97%. Data were analyzed with BD FACS Diva software.

### Removing dead cells using MACS

An alternative approach was used to remove dead cells with the Dead Cell Remove Kit (Miltenyi) following product instructions. Briefly, the cell suspension was centrifuged at 300×g for 10 minutes. Cells were resuspend in 100 μL of Dead Cell Removal MicroBeads and incubated for 15 minutes at room temperature. Dead cells were removed using a MACS Separator (Miltenyi).

### Library construction

Cells were suspended in DMEM-10% FBS. The density and viability of the cells were estimated using Countess II FL Automated Cell Counter (ThermoFisher) upon staining with trypan blue 0.2%. About 10,000 cells were used as an input to generate RNA-seq libraries following the 10X Genomics Chromium Next GEM Single Cell 3′ kit V3 protocol without modification. Libraries were sequenced by commercial deep sequencing facilities.

### Data processing

Single Cell RNA-seq data were processed with Cell Ranger pipeline version 4.0.0. (10x Genomics, Pleasanton, CA, USA), which consisted of two main steps. First, to enhance the mapping of the single-cell RNA-seq reads, a reference transcriptome was created by utilizing the Pig reference genome (Sscrofa11.1) and its annotation downloaded from Ensembl (https://www.ensembl.org/Sus_scrofa/Info/Index). The single-cell RNA-seq reads were mapped using STAR aligner (v. 2.5.1b) against the reference transcriptome to detect and count UMIs and the expressed genes.

### Data analysis

The quality of fresh and frozen single-cell samples were assessed using quality control parameters, including the gene counts per cell, UMI counts per cell and mitochondrial gene expression^39,40^. These quality control parameters were calculated as part of the scRNA-seq data analysis procedure using the Seurat R package version 3.2.1^39^. Since cell doublets or multiplets may exhibit an aberrantly high gene and molecule counts, we set a maximum threshold at 4500 genes or 30,000 molecules (Fig S1). Also, we allowed cells up to 10% mitochondrial gene expression (Fig S1).

The sample data sets were merged into one R Data object per experiment for joint cluster analysis. Then, the count data per cell was normalized and log-transformed using the default settings of the “Normalize Data” function in Seurat^39^. Principal component analysis (PCA) was performed using the highly variable genes for each sample. Significant principal components were selected for subsequent cluster analysis. Single-cell clustering was visualized by UMAP plots with default parameters. Cell types of samples were manually and automatically annotated by marker genes with at least 2-fold increasing in individual cell clusters compared to the remaining cells. For the automated cell type annotation, we used CellMatch. We identified the cluster annotation based on evidence-based score by matching the identified potential marker genes with known cell markers in tissue-specific cell taxonomy reference databases^41^.

For RNA-Seq single-cell trajectory analysis^42,43^, we used an algorithm to learn the changes in the gene expression sequence of each cell. Once the algorithm has learned the overall “trajectory” of gene expression changes, we placed each cell at its proper position in the trajectory (line).

Statistically significant differences between gene levels of cell clusters were calculated by the MAST linear model approach, as implemented in the Seurat package. Genes at least 2-fold deregulated were significantly altered if P-values adjusted for multiple testing were less than 0.001 (Bonferroni correction). The overall similarity of fresh and frozen samples’ gene expression profiles was assessed by correlation performed on “pseudo-bulk” expression profiles^44^, generated by summing counts together for all cells within the same sample by aggregate across function in Scatter package^45^. The raw pseudo-bulk count matrices were normalized using edgeR version 3.30.3^46^. Pearson correlation of the fresh and frozen samples was computed using the normalized counts. The differentially expressed pseudo-bulk genes were identified by edgeR.

### Statistical analysis

The data are presented as the mean ± S.D. We used an unpaired t-test to compare two groups and one-way ANOVA to compare more than two groups. P < 0.05 was considered statistically significant. We used GraphPad Prism 6 for Windows 6.01v to perform statistical analysis.

## Results

### Isolating and preserving high quality single pigskin cells

We used the Miltenyi Human Whole Skin Dissociation Kit and the gentleMACS Dissociator to dissociate pig skins. Our results showed the combination efficiently dissociated pig skin into single cell with only a few cell aggregates and no cell or tissue debris. The resultant cells had healthy and spherical morphology (Fig 1a, b). Live/dead cell staining showed majority of cells were live and confirmed cell aggregates were few (Fig 1c, b). With flow cytometry, we confirmed that >60% of cells were viable (Fig 1e).

**Figure 1.**
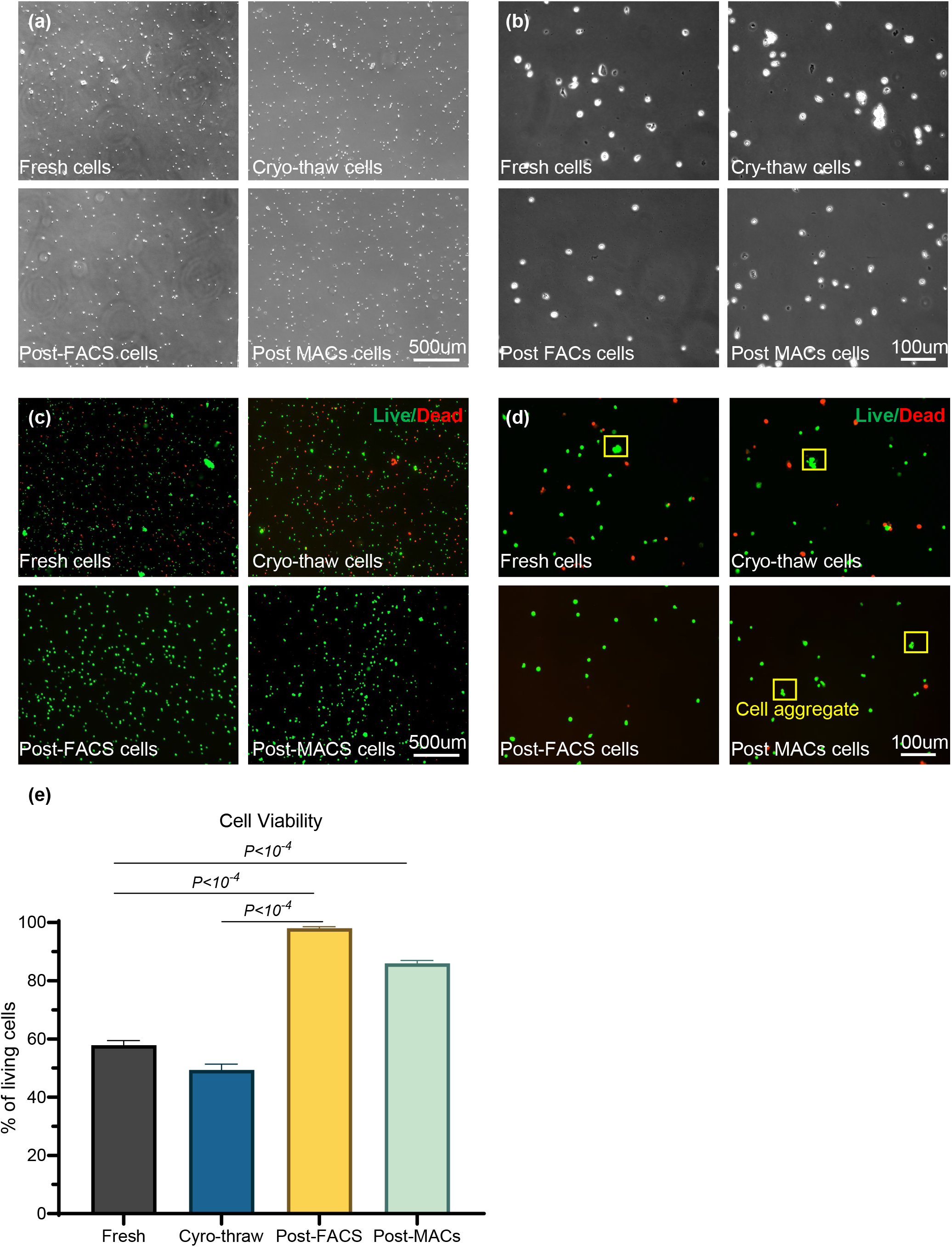
Efficient single cell preparation. Phase images (a, b), live/dead staining (c, d) and flow cytometry viability quantification of fresh isolated, cryopreserved (after thaw), post-FACS and post-MACS pig skin cells. n=3.

We froze the single pig skin cells in 90% FBS + 10% DMSO at −80°C followed by long-term storage in liquid N_2_. After thawing, frozen cells had similar spherical morphology and viability as the fresh cells (Fig 1a-e), preliminarily indicating this method is appropriate for preserving cells. The cryo-preservation did not induce cell aggregation.

Cell aggregates interfere with the downstream library preparation. RNAs released from dead cells negatively affect scRNA-Seq results. We sought to remove both aggregated and dead cells with FACS before library preparation. We used ethidium homodimer-1 to stain dead cells, and side-scatter and forward-scatter profiles to eliminate cell doublets. FACS quantitively removed both (Fig 1d). Since many researchers have difficulty to assess FACS, we also used an alternative and compact device, the Miltenyi MACS, to remove the dead cells. MACS quantitively removed dead cells, however, it was not efficient to remove cell aggregates (Fig 1d). In summary, we suggest that it is appropriated use the Human Whole Skin Dissociation Kit, the gentleMACS Dissociator, DMSO-based cryopreservation and FACS to obtain high quality single cells for scRNA-Seq.

### DMSO-based cryopreservation preserved gene expression

Our data showed the mean genes per cells, mean molecules per cells, mean % mitochondrial genes and their distribution were similar between the fresh and the frozen sample (Fig 2a-c, Table 1). Also, we identified similar distribution when compared fresh and frozen samples by individual cell type (Fig 2a-c). Our data suggest that the DMSO-based cryopreservation approach resulted in high quality scRNA library with minimal transcriptomic alterations (Fig. 3 b, d).

**Figure 2.**
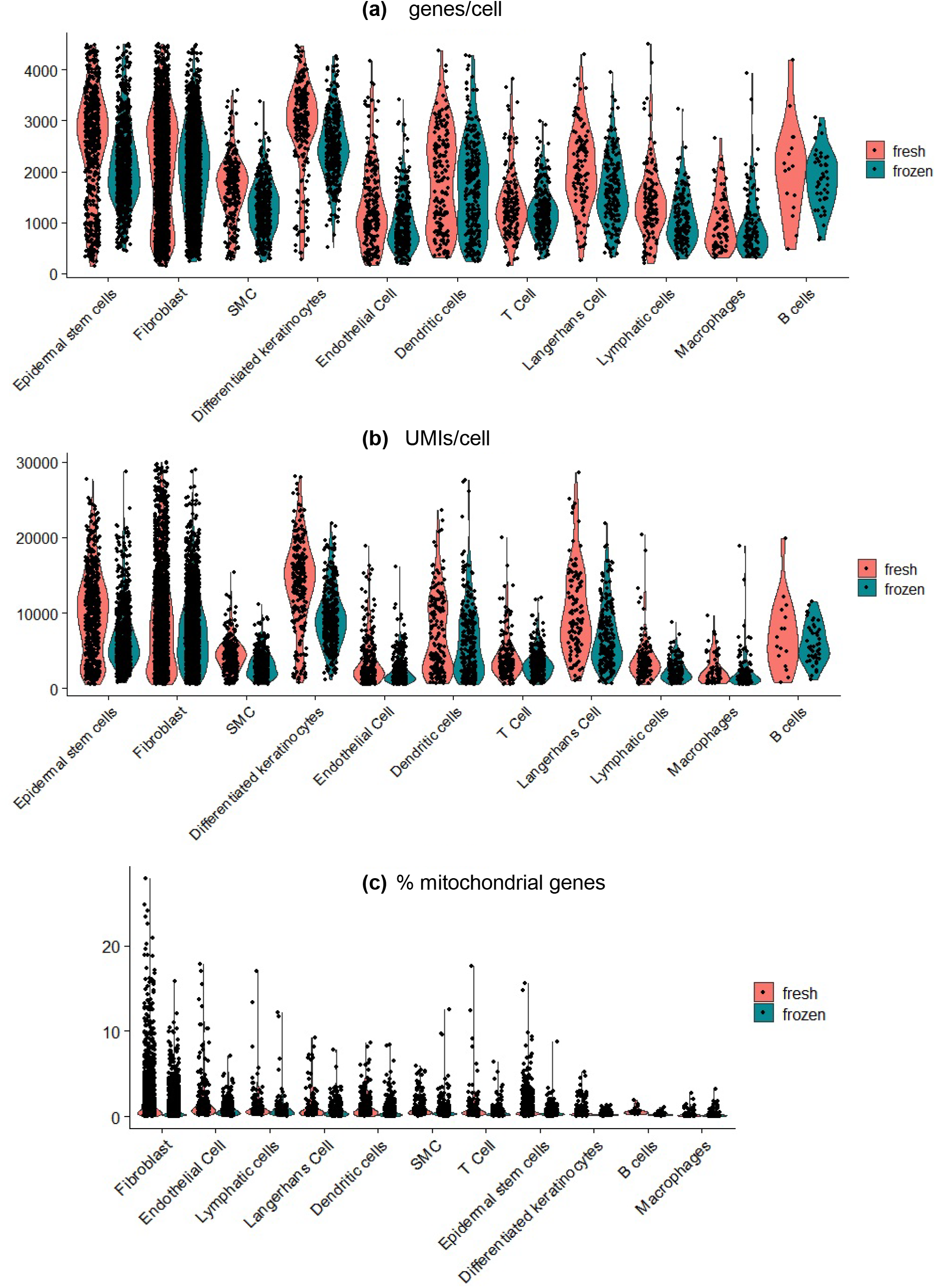
Fresh and frozen cells have similar quality control parameters. The genes/cell (a), UMI/cell (b) and % mitochondrial genes (c) in each cell of fresh and frozen sample by cell type. The distribution and mean are similar between fresh and frozen samples. Each dot represents one cell.

**Table 1:** Cellular composition in fresh and frozen sample.

|  | % frozen | % fresh |
| --- | --- | --- |
| Epidermal stem cells | 17.80 | 16.05 |
| Fibroblast | 37.99 | 45.45 |
| SMC | 8.28 | 7.21 |
| Differentiated keratinocytes | 7.71 | 6.73 |
| Endothelial Cell | 6.29 | 6.54 |
| Dendritic cells | 5.86 | 4.66 |
| T Cell | 5.53 | 4.35 |
| Langerhans Cell | 4.14 | 3.11 |
| Lymphatic cells | 2.94 | 3.37 |
| Macrophages | 2.50 | 2.08 |
| B cells | 0.96 | 0.45 |

**Figure 3.**
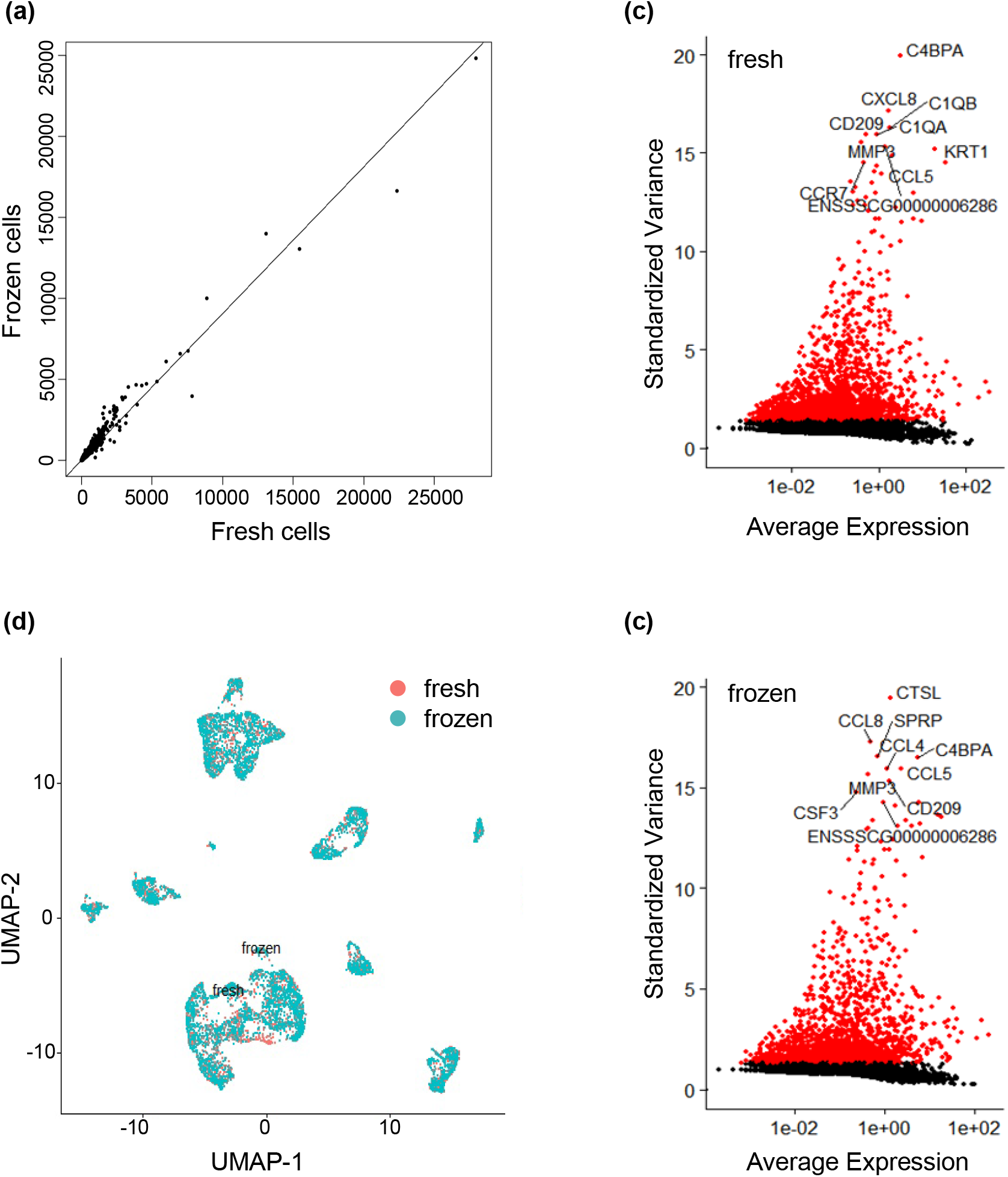
Fresh and frozen cells have similar gene expression profiles. (a) The pseudo-bulk expression profiles of fresh and frozen are compared using correlation scatter plots. The pseudo-bulk expression profile of frozen sample correlate very well with those of fresh cells (R = 0.981). (b, c) UAMP shows fresh and frozen samples have similar structures. (d) Analysis of the subset of genes that exhibit high cell-to-cell variation shows fresh and Frozen samples retain and share the highly variable features with similar variance and average expression.

To assess if cryo-preservation alter the gene expression, the pseudo-bulk expression profiles of frozen and fresh cells were compared using correlation analysis. Pseudo-bulk expression profile of frozen sample correlated very well with those of fresh cells (R = 0.981) (Fig 3a). Among 20,428 expressed genes, there were only 57 differentially expressed genes (DEGs) with between fresh and frozen samples with FDR adjusted P-values < 0.01. This indicates the gene expression profiles between fresh and frozen samples are almost the same. We analyzed the subset of genes that exhibited high cell-to-cell variation in the samples (Fig 3b). Fresh and Frozen samples retained and shared the highly variable features (e.g., CD209, MMP3, CCL5, C4BPA) with similar variance and average expression. Lastly, we assembled the fresh and the frozen sample into an integrated reference, and visualized the integration using a non-linear dimensional reduction UMAP (Fig 3c). The fresh and frozen cells formed similar structures. In summary, these results show the DMSO-based cryopreservation did not alter the gene expression.

### DMSO-cryopreservation approach retained the major skin cell types

Next, we sought to answer if the single cell preparation protocol could retain the major skin cell types and if the DMSO-cryopreservation changed the cell composition. We performed a clustering analysis of the integrated frozen and fresh cells using a graph-based clustering approach^39,47^– the Louvain algorithm^48^. This method embeds cells in a graph structure, clusters cells with similar feature expression, and partition the graph into highly interconnected communities. We identified 18 clusters (Fig 4). Using heatmap, we identified the top 10 marker genes for each cluster (Fig S2). Then, we used a mixed strategy to annotate the cluster identities. Using CellMatch, we annotated clusters based on evidence-based score by matching the identified potential marker genes with known cell markers in tissue-specific cell taxonomy reference databases^48^. Also, we used cell marker genes from recent scRNA-seq studies of skin^17,28^- to identify the remining cell types. This allows us to identify the major skin cell types (Fig 5a). Using heatmap, we identified the top 10 marker genes for each cell type (Fig S3)

**Figure 4.**
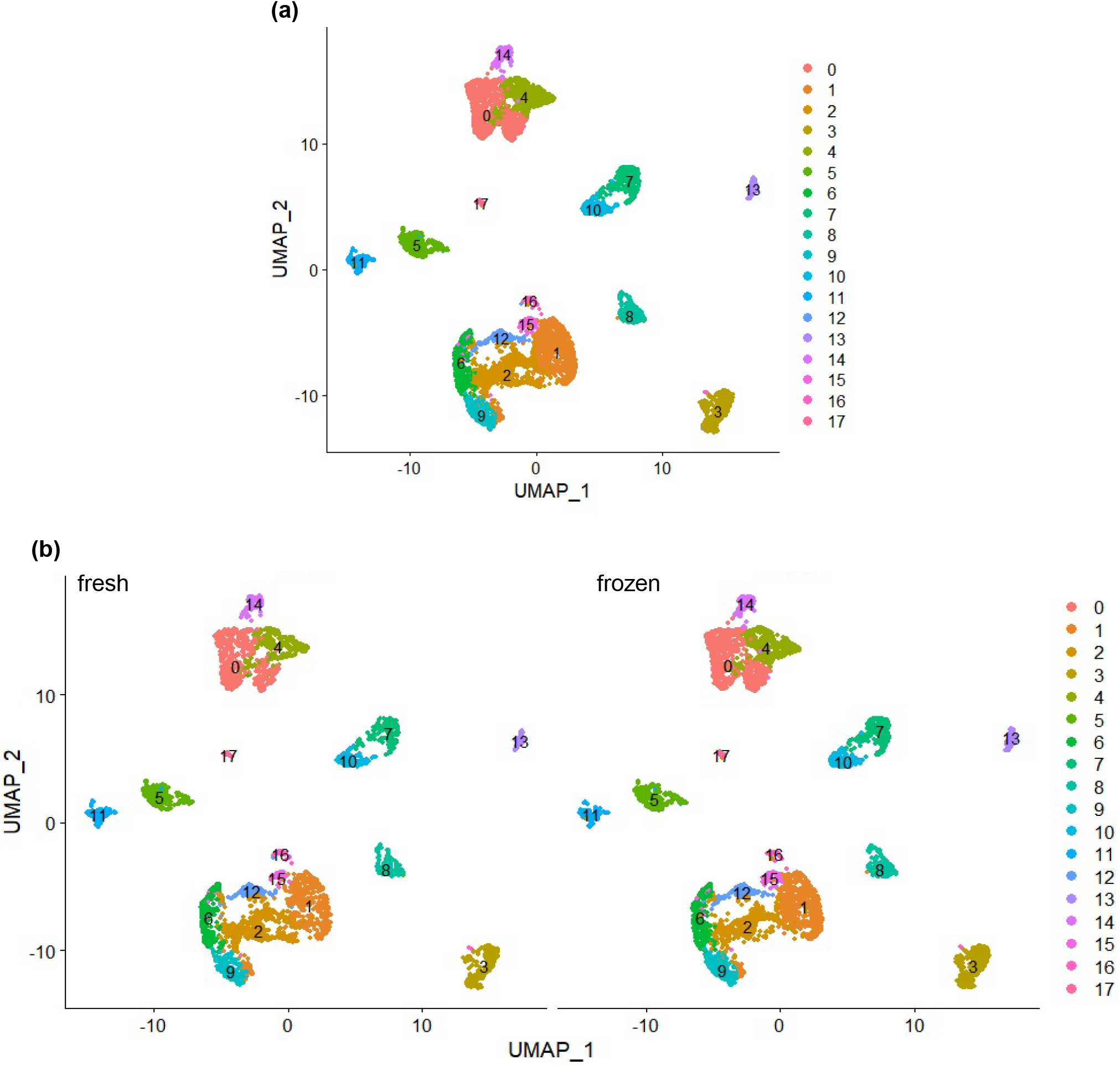
Clustering analysis of combined fresh and frozen cells (a). Fresh and frozen samples had similar clusters.

**Figure 5.**
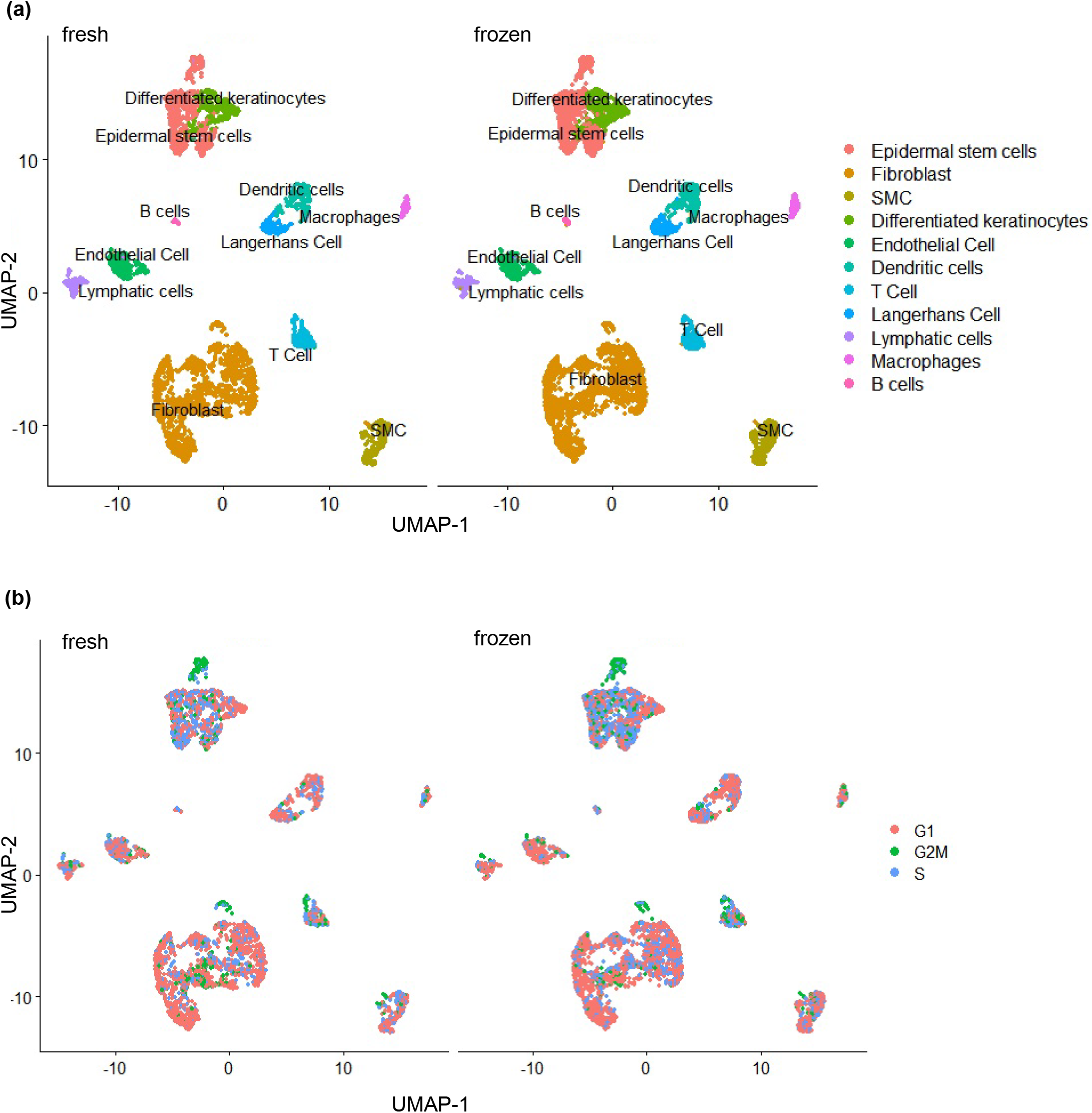
Fresh and frozen samples had similar cell types (a) and cell cycle status (b).

We compared the fresh and the frozen sample in terms of graph structure, cluster, interconnected communities both by cluster (Fig 4b) and by cell type (Fig 5a). The two samples were very similar. Also, DMSO-based cryopreservation kept similar percentage between frozen and fresh cell types (Table 1)

### DMSO-based cryopreservation retained the cell cycle and differentiation trajectory of pig epidermal keratinocytes

Further, we used cell cycle regression to analyze the effect of cryopreservation on the keratinocytes cell cycle. Using this analysis we identified the cell cycle heterogeneity between scRNA-Seq from frozen and fresh samples by calculating cell cycle phase scores based on canonical markers^41^. Our results showed no differences between fresh and frozen keratinocytes regarding in G2/M-phase genes (Fig 5b). To test if fresh and frozen cells could retain the differentiation process, we analyzed if epidermal stem cells differentiate into differentiated keratinocytes. Using RNA-Seq single-cell trajectory analysis ^42,43^, we found in both fresh and frozen samples, epidermal stem cells trajectory differentiate into differentiated keratinocytes (Fig 6a,b).

**Figure 6.**
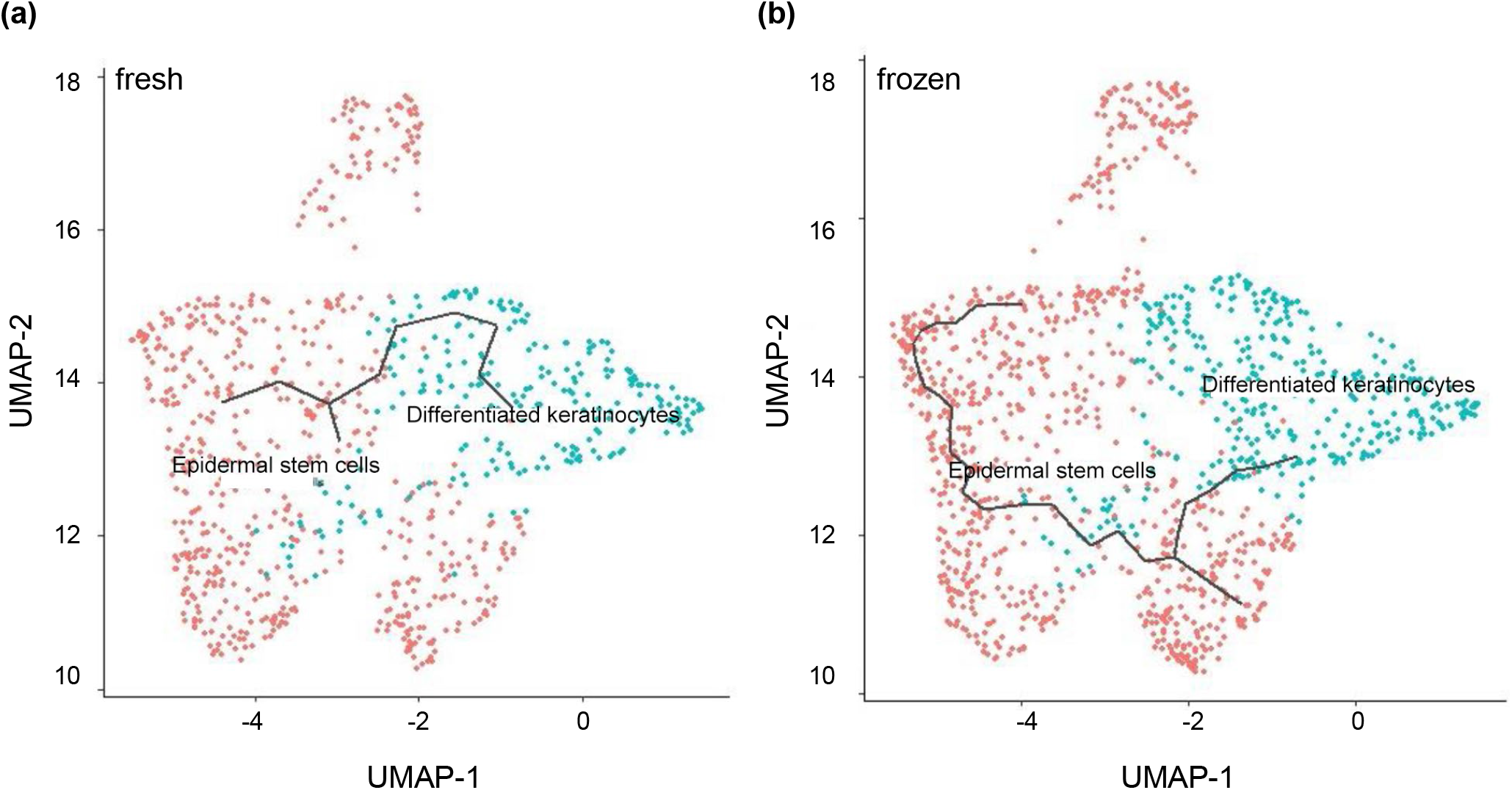
Single-cell trajectory analysis shows epidermal stem cells differentiate into differentiated keratinocytes in both fresh and frozen samples.

## Discussion

Preparing high quality single cells is the first critical step for a successful scRNA-Seq experiment. A good dissociation method should be able to dissolve the whole skin tissue including both epidermis and dermis to release most if not all cells^24,49,50^. The resultant cells should have high viability and few aggregates. A dissociation method with high cell yield and cell viability can minimize the volume of skin sample, and allow detecting cell types with small numbers. To date, a few methods and protocols have been published for preparing skin single cells of rodents^51–57^ and human^58–63^ via enzymatic digestion and mechanical dissociation. These methods diverge in the enzyme types, concentrations, exposure time, temperature^51,52,61–63,53–60^. Additionally, most of them use manual mechanical dissociation, such as dissection, mincing, agitation, pipetting or passing through syringes that are not efficient and consistent. Consequently, the dissociation efficiency, the resultant cell viability, composition are different. This creates challenges when comparing scRNA-Seq results from different labs and using different cell preparation methods. There is a critical need to develop a standardized single cell preparation method. A standard method could be best achieved by using validated commercially available enzyme kits and mechanical dissociation device. Although, we were not able to find commercial kit specifically designed for pig skin, the Miltenyi Human Whole Skin Dissociation Kit and the gentleMACS Dissociator had been used to prepare human skin single cells for scRNA-Seq^24,25,28,29,64,64,65^. Due to the similarity of the human and pig skins, we proposed that the combination of Miltenyi Human Whole Skin Dissociation Kit and gentleMACS Dissociator could also be used to prepare single pig skin cells, which was proven by our results (Fig 1).

With our protocol, the skin tissue was close to completely dissolved. only small numbers of cell aggregates were found, and they could be robustly removed with FACS (Fig 1). We were able to isolate 6.0 −7.5×10^4^ viable cells with a 4-mm full skin biopsy punch. These cells are sufficient for a complete scRNA-seq flow preparing the RNA library cell counting, viability assessment and quality control.

Reliable methods for preserving single cells before scRNA-Seq are thus important and needed. A few methods including using DMSO-based cell culture medium, methanol fixing or commercial cell preservation reagent have been reported to preserve single cells^25,34,36,37^. DMSO aims to minimize forming intracellular big ice crystals, which will damage and kill cells. Methanol fixing works through dehydrating cells to preserve nucleic acids in a collapsed form at high concentrations. Upon rehydration, nucleic acids can be restored to the original form and harvested for library preparation^64,66^. Literature research on scRNA-Seq cell lines showed the methanol-fixing method resulted in high ambient RNA background and a lower gene expression correlation to the fresh un-preserved cells^34^. The same was true for using commercial reagent CellCover reagent^34^. DMSO plus FBS, on the other hands, did not reduce the cell viability and did not alter the cell composition and gene expression significantly^25,34–37,67^. Our result with pig skin cells agreed well with these literature studies using human or rodent primary cells or cell lines (Figs 1, 2, 3, 5, 6).

In summary, we developed a new method to isolate and preserve high quality single cells from pig skin, and successfully obtained high quality scRNA-Seq data. We showed all the major skin cell types could be identified through the scRNA-Seq experiment. We also showed the DMSO-based cryopreservation did not significantly alter gene expression. Limitations with this study include healthy pig skins from young pigs, and no rodent or human DMSO-based cryopreservation comparison.

## Acknowledgments

This research was supported by the University of Nebraska Research Initiative 2018-2019 (YL and WV); the University of Nebraska-Lincoln start-up (YL); the Nebraska DHHS Stem Cell Grant 2019 (YL and WV); the U.S. Army GRANT10824516 (WV); the Pennsylvania State University start-up (YL); and the Department of Defense, USA, W81XWH-BAA-11-1 (WV). This study was financed in part by the Coordenação de Aperfeiçoamento de Pessoal de Nível Superior – Brasil (CAPES) – Finance Code 88882.434714/2019-01 (EPA and LAV). Confocal microscopy imaging was done in the Morrison Microscopy Core Research Facility at the University of Nebraska, Lincoln.

## Competing interests

N/A

---

**Figure s1.**
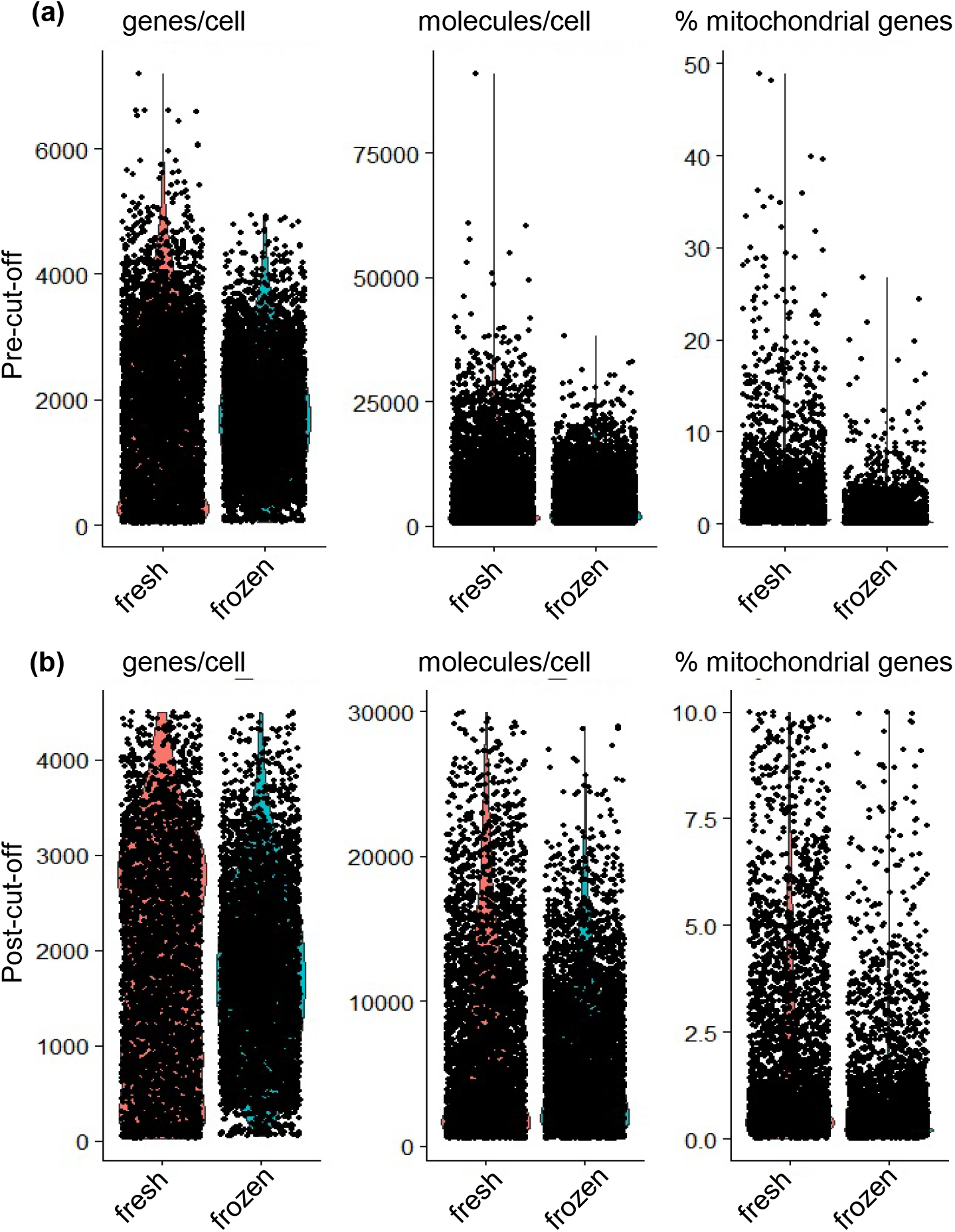
Quality control parameters of fresh and cryo-preserved pig skin cells including number of genes, molecules and % of mitochondrial gene in each cell. (a) pre-cut-off and (b) post-cut-off data are shown. 4500 genes/cell, 30,000 molecules/cell and 10% mitochondrial genes are set as the maximum threshold to exclude the doublets, multiples and low-quality single cells. Each dot represents one cell.

**Figure s2.**
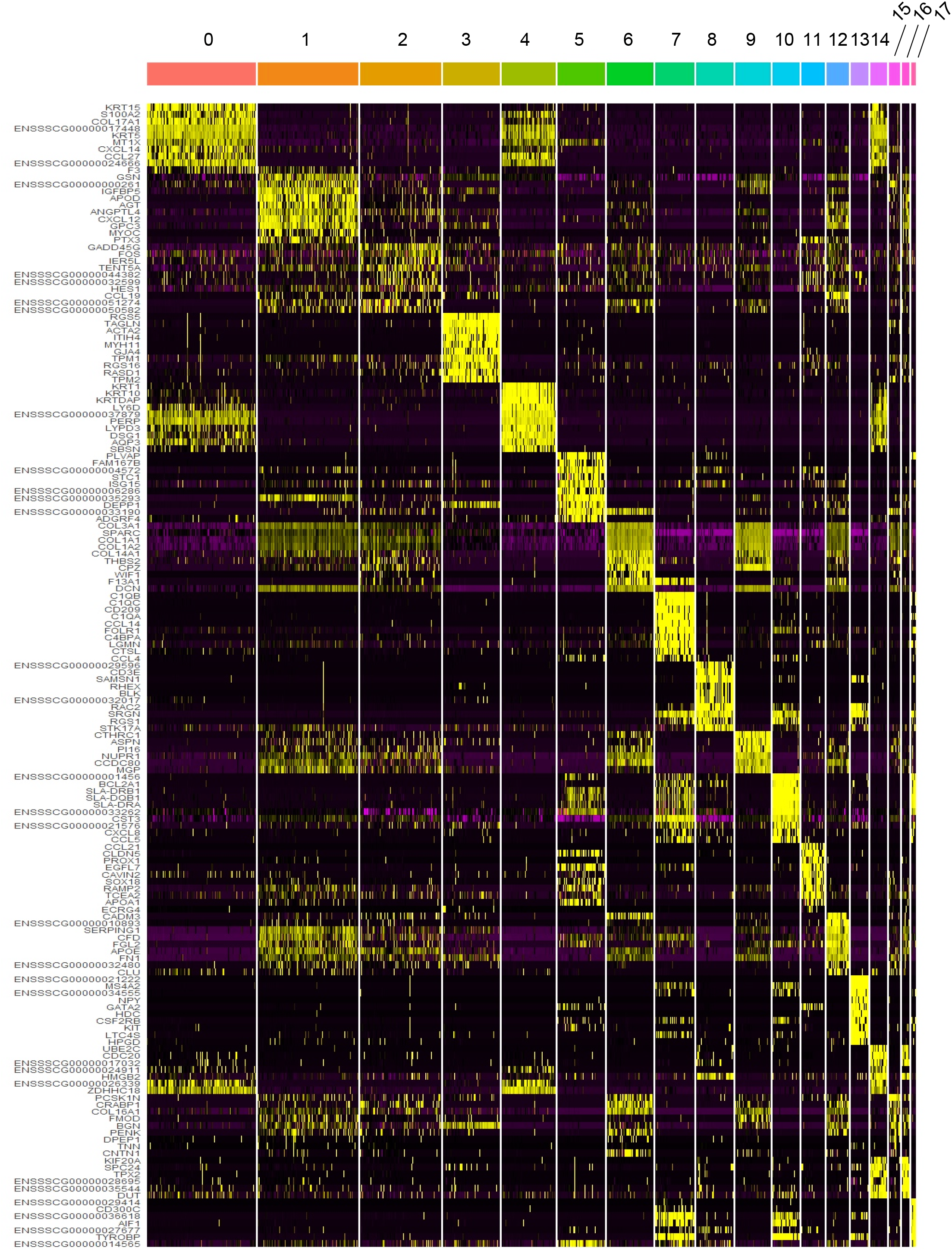
Top 10 marker genes for each cluster. Yellow squares show the differences between one cluster from the other cluster (here numbers from 0 to 17 on the top).

**Figure s3.**
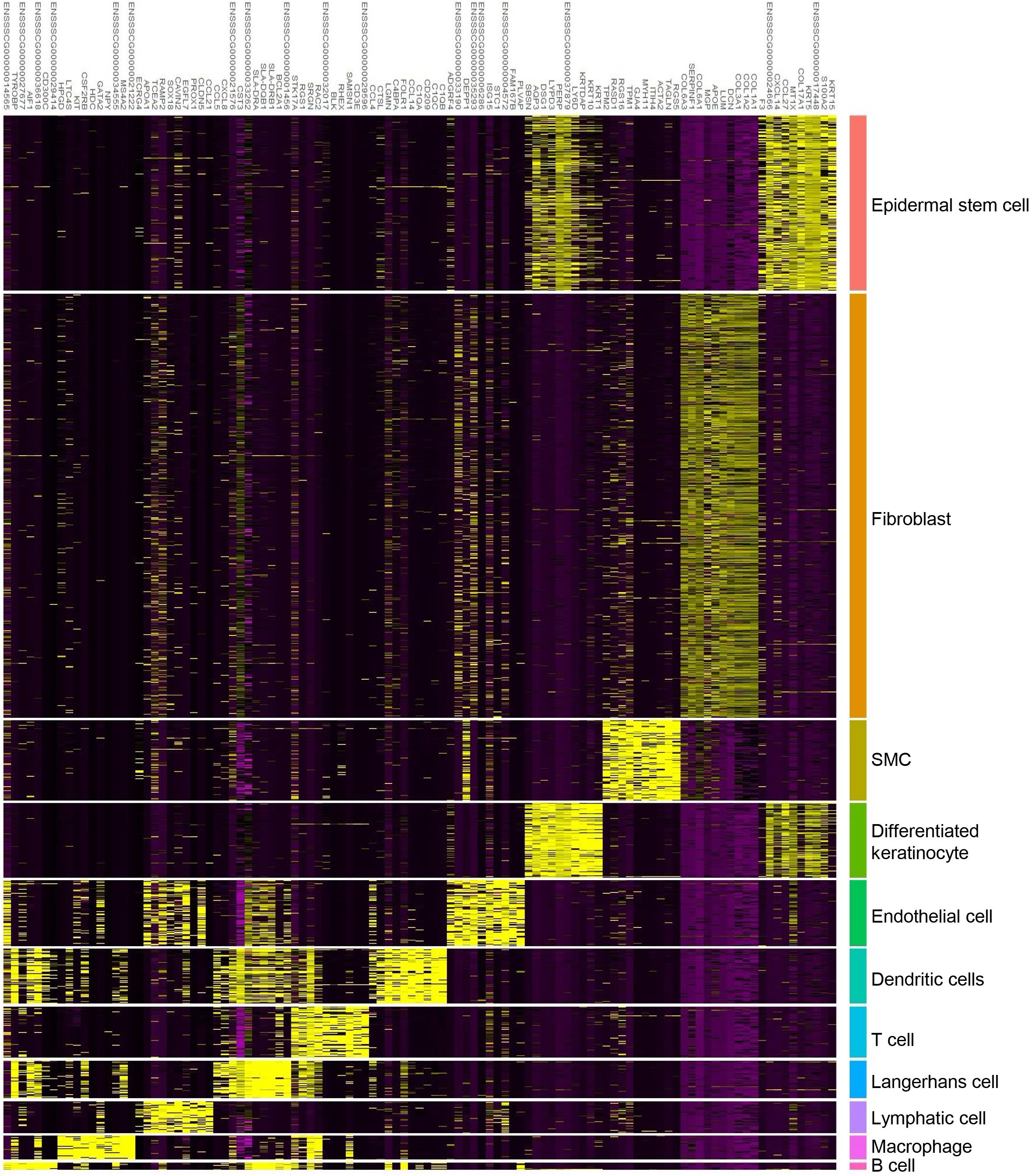
Top 10 marker genes for each cell type. Yellow squares show the differences between cell types.

## Notes

### Competing Interest Statement

The authors have declared no competing interest.

